# Changes in lipid and proteome composition accompany growth of *Bacillus subterraneus* MITOT1 under supercritical CO_2_ and may promote acclimation to associated stresses

**DOI:** 10.1101/320424

**Authors:** Kyle C. Peet, Kodihalli C. Ravindra, John S. Wishnok, Roger E. Summons, Janelle R. Thompson

**Affiliations:** Department of Civil and Environmental Engineering, Massachusetts Institute of Technology, Cambridge, MA, 02139; Department of Biological Engineering, Massachusetts Institute of Technology, Cambridge, MA, 02139; Department of Earth, Atmospheric and Planetary Sciences, Massachusetts Institute of Technology, Cambridge, MA02139

**Keywords:** CO_2_ Sequestration, GCS, supercritical CO_2_, Bacillus, biocompatibility, Proteomics

## Abstract

Recent demonstration that multiple *Bacillus* strains grow in batch bioreactors containing supercritical (sc) CO_2_ (i.e. >73 atm, >31°C) is surprising given the recognized roles of scCO_2_ as a sterilant and solvent. Growth under scCO_2_ is of interest for biotechnological applications and for microbially-enhanced geologic carbon sequestration. We hypothesize that *Bacillus* spp. may alter cell wall and membrane composition in response to scCO_2_-associated stresses. In this study, protein expression and membrane lipids of *B. subterraneus* MITOT1 were profiled in cultures grown under headspaces of 1 and 100 atm of CO_2_ or N_2_. Growth under 100 atm CO_2_ revealed significantly decreased fatty acid branching and increased fatty acyl chain lengths relative to 1 atm cultures. Proteomes of MITOT1 grown under 1 and 100 atm pressures of CO_2_ and N_2_ were similar (Spearman R>0.65), and principal component analysis revealed variation by treatment with the first two principal components corresponding to headspace gas (CO_2_ or N_2_) and pressure (1 atm and 100 atm), respectively. Amino acid metabolic proteins were enriched under CO_2_, including the glycine cleavage system, previously shown to be upregulated in acid stress response. These results provide insights into the stationary phase physiology of strains grown under scCO_2_, suggesting modifications of cell membranes and amino acid metabolism may be involved in response to acidic, high CO_2_ conditions under scCO_2_.

## Introduction

Supercritical (sc) phase CO_2_ is employed as an industrial solvent and sterilizing agent, but the bactericidal properties of scCO_2_ have been a limiting factor for development of two phase systems for biological growth coupled to in situ extraction (1) and for harnessing microbial processes during geologic carbon sequestration (GCS) (2). However, recent demonstration of microbial growth under scCO_2_ (3) opens new prospects in biotechnology and GCS applications where scCO_2_ is present, while challenging the efficacy of scCO_2_ sterilization in food and medical industries.

ScCO_2_ exposure presents a complex set of stresses for cells due to a combination of factors including cytoplasm acidification, increased dissolved CO_2_ and CO_2_ anion concentration, cell membrane permeabilization, extraction of non-polar molecules into a scCO_2_ phase, and physical cell rupture (4–11). Previous studies examining responses of bacteria to scCO_2_ exposure have relied on short duration exposures (less than 1 hour) and revealed changes to lipid acyl chains (11, 12) and to lipid head groups, with a reduction in phosphatidylglycerol lipids (11) and increased production of leucine and isoleucine (13), suggesting these amino acids may be involved in a short term scCO_2_ stress response. In all cases, previous studies of bacterial response to short term scCO_2_ exposure lead to cell death, providing little time for cells to acclimate before succumbing to lethal stresses associated with scCO_2_ exposure. However these studies do not represent profiles of cells capable of acclimation and growth under scCO_2_. Nevertheless, the cellular responses observed during these experiments suggest that alterations to cell membranes and global changes to protein expression may be necessary to acclimate to scCO_2_-assocated stresses.

We have recently documented growth of *Bacillus* spp. under a scCO_2_ headspace (3), however how cells acclimate to grow under scCO_2_ is unknown. Individual factors associated with scCO_2_ (e.g. elevated pressure, solvent, and acid stress) may have opposing influences on cellular acclimation. High pressures compress membrane lipids with lipid disordering caused by dissolution of gases into membrane bilayers (14). Bacteria under high pressure may compensate by producing more unsaturated lipids or by decreasing lipid chain length in order to increase and maintain membrane fluidity (15, 16), a response similar to that observed following exposure to low temperature (16–20). However, solvent stress has been demonstrated to increase the proportion of saturated fatty acid content and increasing acyl chain lengths (21). Bacteria also alter membrane lipids in response to pH and oxygen availability. Acid stress in *B. subtilis* and *Clostridium acetobutylicum* was linked to a decreased proportion of branched and unsaturated fatty acids, which was suggested to increase membrane rigidity and decrease the proton flux across membranes (22, 23). Anaerobic growth in *B. subtilis* results in increased lipid chain length (24) which may also promote membrane rigidity.

Since the effects of pressure may have opposing effects on membrane fluidity when compared with solvent and acid stress, it is not yet apparent how cells exposed to scCO_2_ will balance production of more fluid membrane lipids expected under high pressure with more rigid membrane lipids associated with solvent or acid-stress. We hypothesize that cells acclimated to scCO_2_ may display a lipid profile that is intermediate between acid/ solvent stressed and pressure stressed phenotypes. Alternatively, they may have a profile more similar to one of those conditions if one aspect of scCO_2_ is a more severe stress.

Multiple studies have examined responses of protein or gene expression to individual stresses associated with scCO_2_ (e.g. low pH, high pressure, high CO_2_ concentration), but not necessarily the combined stress of high pressure (>73 atm), low pH (<4), and high CO_2_ concentration (>2.5 M)(3). Such studies suggest that mechanisms for acclimation to low pH include upregulation of proton-pumping ATP transporters (25), transport/metabolism of amino acids that buffer intracellular pH (26, 27), expression of amino acid decarboxylating enzymes through consumption of intracellular protons (28), while various enzymes (e.g. urease and the arginine deiminase system) can produce alkaline products from amino acids and other compounds to buffer intracellular pH (29–32). Responses of cells acclimated to growth at 1 atm to elevated pressures share similarities with a profile of general stress response, e.g. upregulation of sigma factors and chaperone proteins, and reduced biomass and protein expression (33–36) and may also include shifts in expression of various housekeeping functions such as transcription, translation and metabolism (33, 34). In contrast, in barophilic organisms where high pressures (e.g. 280-700 atm) are optimal growth conditions, exposure to pressure is associated with upregulation of specific respiratory genes (37, 38), and down-regulation of chaperone proteins (39).

To better understand how bacteria may acclimate to grow under scCO_2_, we have examined how the lipid content and proteome of *B. subterraneus* MITOT1 varies in stationary phase cultures as a function of anaerobic N_2_ and CO_2_ headspaces at ambient pressure (1 atm) or at 100 atm pressure. *B. subterraneus* MITOT1 was enriched and isolated from a deep subsurface rock core from the Paaratte Formation, Otway Basin in Southern Australia (3, 40). This strain was among several *Bacillus* spp. shown to germinate from spores and grow into vegetative cells under scCO_2_; where physiological characterization suggests it is a facultative aerobe with optimal growth at 1 atm of pressure (3). Results from proteome and lipid analyses suggest several modifications of lipid and proteome profiles that may facilitate growth under scCO_2_, but also point to a high degree of similarity across all anaerobic conditions, suggesting the homeostasis of stationary phase cultures is maintained by similar mechanisms independent of responses to scCO_2_ exposure. This study presents the first examination of acclimation responses to bacteria grown under an environment containing scCO_2_.

## Methods

### Chemicals

DL-Dithiothreitol (CAS: 3483-12-3), iodoacetamide (CAS: 144-48-9), acetonitrile (CAS: 75-05-8), formic acid (CAS: 64-18-6) were purchased from Sigma-Aldrich. Sequencing grade modified trypsin (Cat. no. V5111) and ProteaseMAX™ Surfactant (Cat. no. V2071) were purchased from Promega. Imm. Drystrip pH 3-11 (Cat No. 17600377) and buffer (Cat No. 17600440) purchased from GE. OMIX tips are from Agilent technologies (Part No. A57003100). VIVASPIN 500 (Cat. no. VS0191) 3K molecular weight cut-off filters are purchased from Sartorius. Distilled water was prepared in-house with double distillation. Unless otherwise noted, all the materials were obtained from commercially available sources and were used without further purification.

### Cell growth and collection for lipid and protein analyses

All cultures were started from spore inocula of 10^5^/ml (direct counts). Spores were prepared as described previously (3). Growth media for *B. subterraneus* MITOT1 consisted of MS medium: (in g/l) 0.5 yeast extract, 0.5 tryptic peptone, 10.0 NaCl, 1.0 NH_4_Cl, 1.0 MgCl_2_.6H_2_O, 0.4 K_2_HPO_4_, 0.4 CaCl_2_, 0.0025 EDTA, 0.00025 CoCl_2_.6H_2_O, 0.0005 MnCl_2_.4H_2_O, 0.0005 FeSO_4_.7H_2_O, 0.0005 ZnCl_2_, 0.0002 AlCl_3_.6H_2_O, 0.00015 Na_2_WoO_4_.2H_2_O, 0.0001 NiSO_4_.6H_2_O, 0.00005 H_2_SeO_3_, 0.00005 H_3_BO_3_, and 0.00005 NaMoO_4_.2H_2_O. With a supplement for metal reducers consisting of 1.3 g/l MnO_2_, 2.14 g/l Fe(OH)_3_, and 1.64 g/l sodium acetate (41). All culture media for anaerobic conditions was degassed with the respective N_2_ / CO_2_ headspace for 30 minutes, followed by addition of Na_2_S in an anaerobic chamber to further purge any residual oxygen. After completion of incubation, high-pressure samples were depressurized at a rate of 3-5 atm^-1^ minute over approximately 30 minutes. Cells were collected by centrifugation at 14,000 X g for 6 minutes, followed by resuspension in sterile filtered PBS and re-centrifugation. Cell pellets were frozen in -80 °C until further analysis. Aliquots of biomass were taken for direct cell counts and viable cell counts; briefly, total cells were observed by Syto9 staining and epifluorescent microcopy and viable cells were enumerated by cell plating on LB agar, as described previously in (3). The pH of culture media under various headspaces and pressures was measured by visualization of a pH indicator strip (EMD Chemicals) through a 25-ml view cell (Thar Technologies; 05422-2). Media was pH 7 for 1 and 100 atm N_2_ headspaces, and pH 5 and 3.5 for 1 and 100 atm CO_2_ headspaces, respectively.

Aerobic cultures of MITOT1 were incubated for 72 hours with shaking at 37°C to target late stationary phase (OD > 5), before sporulation occurred. Anaerobic cultures of MITOT1 were grown in 316 stainless steel vessels as described in Peet et al. (3) in triplicate, shaking at 37 °C in the following four conditions: 1) 1 atm, 100% N_2_ headspace; 2) 1 atm, 95% CO_2_, 5% H_2_ headspace (referred to as 1 atm CO_2_); 3) 100 atm, 100% N_2_ headspace; 4) 100 atm, 97% CO_2_, 3% He headspace (referred to as 100 atm CO_2_). Sampling times of 21 and 30 days for MITOT1 cultures grown under 1 atm and 100 atm, respectively, were selected to target similar durations of time spent in stationary phase (>7 days) in order to minimize variability associated with growth phase in cultures. Sampling times were estimated based on observed dynamics under 1 atm (Supplemental Fig. 1) and the conditional probability of growth under 100 atm (Supplemental Fig. 2) based on application of Bayes theorem to a logistic regression model for growth outcome (observed/not observed) for MITOT1 as a function of time with an inocula of 10^5^ spores per ml (3). Final cell densities of reactors with positive growth under the four conditions were measured by direct counts and compared by ANOVA.

### Lipid extraction and construction of fatty acid methyl esters (FAMEs)

Triplicate samples for aerobic stationary phase cultures and each of the four anaerobic growth conditions were extracted by a modified Bligh-Dyer extraction for polar lipids (42). All vials, glass pipettes and foil were combusted before use. Syringes were triple washed in each of the following solvents before and between pipetting: methanol, dichloromethane (DCM), and hexane. 1 ml of a solvent mixture containing methanol: DCM: phosphate buffered saline (PBS) (10:5:4) was added to the 1.5 ml centrifuge tube containing frozen biomass and the pellet was resuspended and transferred to a 50 ml glass centrifuge tube. The original 1.5 ml centrifuge tube was washed twice more with this solvent mixture to collect all cells, and a total of approximately 7 ml of solvent was added to the 50 ml glass tube. 1 µg of 3-Methylheneicosane was added as an internal standard to each sample at the beginning of extractions. Cells in this solvent mixture were vortexed for 5 minutes, followed by 15 minutes of sonication to further extract lipids and then centrifuged for 10 minutes at 720 X g. Approximately 90% of the solvent mixture was aspirated to a new glass vial without disturbing cells, and this extraction process was repeated once with the same solvent mixture, twice with a solvent mixture containing methanol: DCM: 2.5 % trichloroacetic acid (10:5:4), and once with DCM: methanol (3:1), pooling all the extractions in a separate vial. Phase separation was conducted by adding 5 ml of PBS and 5 ml of DCM with gentle shaking, followed by removal of the lower (DCM) phase to a new vial. 5 ml of DCM was added twice more to re-extract aqueous phase, and the pooled DCM phases were evaporated in a Turbovap at 37 °C under a stream of 100% ultrapure N_2_. The concentrated samples (of approximately 1 ml volume) were transferred to 4 ml vials and then dried down completely. These were labeled as total lipid extracts (TLE) and stored at 4 °C. Intact polar diacylglycerols were converted to FAMEs by methanolysis. Briefly, TLE’s were resuspended in 200 µl of DCM: methanol (9:1), removing 140 µl to a 2 ml vial with insert, and then drying down the 140 µl. Once dried, 100 µl of methanoic HCl was added, samples were capped and heated at 60°C for greater than 1.5 hours. Samples were evaporated, followed by addition of 200 µl of DCM: methanol (9:1), evaporation, addition of 200 µl of methanol, evaporation, and then resuspension in hexane for analysis via GC/MS.

### Determination of unsaturated double-bond positions

Monounsaturated fatty acid double-bond positions were determined by the method of Yamamoto et al. (43). An aliquot of FAMEs for each sample with detectable unsaturated fatty acids based on GC/MS analysis of TLE’s was transferred to a new 4 ml vial and dried down. 100 µL of dimethyl disulfide (DMDS) and 50 µL of iodine was added to each vial and vials were heated on a heating block at 35°C for 30 minutes. 1 ml of hexane was added to each vial and Na_2_S_2_O_3_ was added to vials 20 µl at a time until mixture turned clear, with vigorous vortexing between each addition. The organic layer was removed and then re-extracted with DCM: methanol (9:1). Organic layers were combined and run through a Na_2_SO_4_ column to remove residual water, followed by rinsing the column with DCM and hexane, before concentrating under N_2_ gas. Samples were resuspended in hexane and analyzed via GC/MS.

### Analysis of lipids

Samples were analyzed on an Agilent 7890A gas chromatograph attached to an Agilent 5975C mass selective detector (MSD) equipped with a programmable temperature vaporization (PTV) injector. 1 µl of sample dissolved in hexane was injected into an Agilent J&W DB-5ms column (60 meter X 0.25 mm internal diameter, with 0.25 µm coating). The column was held at 60°C for 2 minutes following injection, then ramped up to 150°C at 10°C per minute, followed by ramping up to 315°C at 3°C per minute. Total run time was 90 minutes per sample. GC/MS peaks and spectra were acquired with Agilent GC/MSD software and peak areas were manually integrated with Enhanced Data Analysis software. Lipids were identified by searching the mass spectra of integrated peaks against the National Institute of Standards and Technology (NIST) database and matching retention times and peak elution order to NIST predictions. To search for hopanoids, m/z 191 was used to indicate the potential presence of hopanoids (44). Ion 191 was extracted from chromatograms and any spectra with large 191 ions were searched against the NIST database to compare mass spectra. Lipids were normalized to the internal standard for comparison between samples. The normalized means of triplicate extraction blanks were subtracted from all n16:0 and n18:0 peak areas in all samples. The uncertainty from extraction blanks was propagated into error bars shown in all figures. Average acyl chain length was calculated by multiplying each fatty acid’s fractional abundance by the length of the acyl chain and summing those weighted values. Microsoft Excel was used for T-tests and analysis of variance (ANOVA) was calculated with JMP Pro 11 (SAS software).

### Protein extraction and purification

Following growth of cultures, cells were collected by centrifugation (14,000 X g for 6 min), washed in PBS and frozen at -80°C until extraction. Whole cell proteomes were extracted by adding 200 µl of lysis buffer (Huang et al. 2012) containing 8 M urea, 4% (w/v) CHAPS, 40 mM Tris and 65 mM DTT to each frozen cell pellet. 100 µl of sterile 0.1 mm zirconia beads was added to each tube and samples were bead beat for 1 min at maximum speed (MOBio vortexer), followed by 30 seconds on ice. Bead beating and ice bath cooling was repeated 10 times, with the final removal of beads and cell debris by centrifugation for 3 min at 14,000 X g. The protein-containing supernatant was aspirated, placed in a new tube, and frozen at -80°C until digestion. An aliquot of each sample was used for protein quantification via BioRad Protein Assay with bovine serum albumin standard.

Proteins were purified using optimized conditions for proteome analysis. Cell lysates were added to Vivaspin columns (Sartorius) with a 3000 Da size cutoff to remove urea and extraction buffer reagents. An additional 500 µl of water was added to each column to dilute the sample and columns were spun for 10 min at 13,000 X g, room temperature. This was repeated, with the flow-through discarded each time. The remaining proteins in the column were resuspended and added to a new Eppendorf tube with four volumes of cold acetone (-20°C). Proteins were precipitated by incubating tubes at -80°C for 30 minutes and then centrifuging at 16,000 X g for 10 minutes at 4°C. The supernatant was discarded and the protein pellet was washed with 300 µl acetone and spun again, followed by air drying for 5 minutes to allow residual acetone to evaporate.

One set of replicates (Batch 1), with 1 sample from each condition, was processed in the manner described above, but with an additional initial step for precipitation of proteins with acetone prior to application to the Vivaspin columns in order to remove urea and extraction buffer reagents. This initial acetone precipitation step was associated with product loss during the first batch of extractions, and was deemed unnecessary for processing a second batch of proteomes with 2 samples from each conditions. Batch 1 samples are noted in figures and tables and are included for qualitative or univariate comparison to other replicates.

### Protein digestion and peptide fractionation

Proteins were re-suspended in a 15 µl of 8M urea (dissolved in 50 mM ammonium bicarbonate) followed by adding 20 µl of 0.2% ProteaseMAX^TM^ (Promega) surfactant, 50 µl of ammonium bicarbonate, and 2.12 µl of 400 mM dithiothreitol (DTT) to reduce disulfide bonds. Samples were incubated at 56°C for 30 minutes, and then alkylated by addition of 6 µl of 550 mM iodoacetamide, followed by incubation for 30 minutes at room temperature in the dark. To prevent alkylation of trypsin, excess iodoacetamide was inactivated by addition of 2.12 µl of DTT and incubated for an additional 30 minutes in the dark. Proteins were digested by adding 3.7 µl of 0.5 µg/µl trypsin (1:27 trypsin: protein ratio) and 1 µl of 1% ProteaseMAX^TM^ followed by 3 hours incubation at 37°C. After digestion, trypsin was inactivated by addition of 20% trifluoroacetic acid to a final concentration of 0.5%. Digested proteins were concentrated and desalted with OMIX tips (Agilent technologies, Part No. A57003100) according to manufacturer instructions, and dehydrated to dryness in a SpeedVac^®^.

To fractionate peptides by isoelectric point, samples were re-suspended in 3.6 ml of 1X off-gel buffer and then loaded onto an Agilent off-gel fractionator with IPG strips (pH 3-11) according to manufacturer instructions. For the first 4 samples, the 24 fractions were pooled into 19 fractions (combining 1 and 24, 2 and 23, 3 and 22, 4 and 21, 5 and 20, 6 and 19, without combining fractions 7-18). As protein concentrations were relatively low and LC-MS runs did not show a large number of peptides, we pooled the 24 fractions down to 12 fractions in the second set of 8 samples. All fractions were dried in a SpeedVac^®^ prior to re-suspension in 20 µl of 98% H_2_O, 2% acetonitrile, and 0.1% formic acid for LC-MS analysis as described below.

### LC-MS parameters and protein profiling

An Agilent 6530 quadrupole time-of-flight (QTOF) mass spectrometer equipped with an electrospray ionization (ESI) source was used. All samples were analyzed using an Agilent 1290 series Ultra Performance Liquid Chromatography (UPLC) (Agilent Technologies, Santa Clara, CA, USA) containing a binary pump, degasser, well-plate auto-sampler with thermostat, and thermostatted column compartment. Mass spectra were acquired in the 3200 Da extended dynamic range mode (2 GHz) using the following settings: ESI capillary voltage, 3800 V; fragmentor, 150 V; nebulizer gas, 30 psig; drying gas, 8 L/min; drying temperature, 380°C. Data were acquired at a rate of 6 MS spectra per second and 3 MS/MS spectra per second in the mass ranges of m/z 100–2000 for MS, and 50-2500 for MS/MS and stored in profile mode with a maximum of five precursors per cycle. Fragmentation energy was applied at a slope of 3.0 V/100 Da with a 3.0 offset. Mass accuracy was maintained by continually sprayed internal reference ions, *m/z* 121.0509 and 922.0098, in positive mode. Agilent ZORBAX 300SB-C18 RRHD column 2.1 × 100 mm, 1.8μm (Agilent Technologies, Santa Clara, CA) was used for all analyses. LC parameters: autosampler temperature, 4°C; injection volume, 20 µl; column temperature, 40°C; mobile phases were 0.1% formic acid in water (phase A) and 0.1% formic acid in acetonitrile (phase B). The gradient started at 2% B at 400 µl/min for 1 min, increased to 50% B from 1 to 19 min with a flow rate of 250 µl/min, then increased to 95% B from 19 to 23 min with an increased flow rate of 400 µl/min and held up to 27 min at 95%B before decreasing to 2% B at 27.2, ending at 30 min and followed by a 2 minute post run at 2% B.

### Protein data processing

Raw data was extracted and searched using the Spectrum Mill search engine (B.04.00.127, Agilent Technologies, Palo Alto, CA). “Peak picking” was performed within Spectrum Mill with the following parameters: signal-to-noise was set at 25:1, a maximum charge state of 7 is allowed (z = 7), and the program was directed to attempt to “find” a precursor charge state. During peptide searching the following parameters were applied: peptides were searched against the genome of *B. subterraneus* MITOT1(40), carbamidomethylation was added as a fixed modification, and the digestion was set to trypsin. Additionally, a maximum of 2 missed cleavages, a precursor mass tolerance +/-20 ppm, product mass tolerance +/- 50 ppm, and maximum ambiguous precursor charge = 3 were applied. Data were evaluated and protein identifications were considered significant if the following confidence thresholds were met: protein score > 13, individual peptide scores of at least 10, and Scored Peak Intensity (SPI) of at least 70%. The SPI provides an indication of the percent of the total ion intensity that matches the peptide’s MS/MS spectrum. A reverse (random) database search was simultaneously performed and manual inspection of spectra was used to validate the match of the spectrum to the predicted peptide fragmentation pattern, hence increasing confidence in the identification. Standards were run at the beginning of each day and at the end of a set of analyses for quality control. Protein expression values (spectrum counts) were calculated in Scaffold software with the imported peptide hits from Spectrum Mill. The threshold for including a protein was a minimum of two distinct peptides with 95% confidence. To compare between samples, spectrum counts for each protein were divided by the sum of spectrum counts in respective samples, resulting in protein expression values as a percent of total.

Data analysis and statistics were conducted with Microsoft Excel, JMP Pro11, Simca, and Primer 6 software. The Kruskal-Wallis test was used to determine if individual proteins were differentially expressed under culture conditions. Proteins with 2 or more samples below the detection limit were not considered for significance. Clustering was performed with Spearman rank correlation and Principal Component Analysis (PCA) implemented in Primer 6 software (Plymouth Marine Laboratory), for PCA proteins representing at least 0.5% of the spectral counts for an individual sample were included in the analysis. Partial least squares discriminant analysis (PLS-DA) was used to identify proteins that are differentially represented among the different conditions, and Gene Set Enrichment Analysis (GSEA) (45) was used to determine if pathways (or gene sets) are differentially expressed in response to different conditions. The MITOT1 genome (40) was annotated with the Kyoto encyclopedia of genes and genomes (KEGG) automatic annotation server (KAAS) (46) to obtain KEGG IDs for proteins to be used in conjunction with KEGG gene sets in GSEA. Proteins that could not be annotated with KEGG were excluded from this analysis. Pathway (gene set) size was set to a minimum of 5 proteins, and 1000 permutations were performed. Proteins detected by LC/MS with ‘hypothetical’ RAST annotations (40, 47) were submitted to Phyre2 (48) for additional characterization. Phyre2 annotations are included (in Supplemental Table 4) when predicted with greater than 90% confidence, 20% identity, and 30% coverage.

## Results

### Growth of MITOT1 in bioreactors under different headspace and pressures

We observed growth in 5 of 6 bioreactors containing 1 atm CO_2_ headspace, 3 of 3 bioreactors containing 1 atm N_2_ headspace, 4 of 11 bioreactors containing 100 atm CO_2_, and 7 of 7 bioreactors containing 100 atm N_2_. Growth variability under high pressure CO_2_ is consistent with previous observations and has been discussed in detail elsewhere (Peet et al,(3)). When more than 3 replicates were available for analysis, samples with higher biomass were selected in order to maximize material for protein and lipid analyses. All bioreactors with observed biomass were within a factor of 5 of the maximum cell density observed in previous experiments carried out under anaerobic conditions (i.e. 10^7^ to 2×10^8^ cells/ml) (3), consistent with these cultures being in, or entering, stationary phase (Supplemental Fig. 3). Direct counts varied significantly between headspace and pressure/incubation conditions, with greater counts in CO_2_-incubated cultures (p= 0.005, F-ratio= 15.1), and in high pressure cultures (p= 0.003, F-ratio= 18.1), while the interaction of pressure and headspace gas was not significant. No significant differences in viable counts (cfu/ml) were observed between headspace or pressure/incubation, which could have indicated differences in growth phase (p>0.05; two-factor ANOVA).

### *B. subterraneus* MITOT1 lipids under supercritical CO_2_ appear similar to an acid stressed phenotype

Since previous studies have shown changes in lipid composition associated with pH, pressure and other stresses (15, 17, 18, 22–24, 49), we examined how cultures grown under 1 atm CO_2_, 1 atm N_2_, 100 atm CO_2_, and 100 atm N_2_ differed with respect to lipid composition, branching, saturation, and chain length. Lipids from *B. subterraneus* MITOT1 grown under aerobic conditions with ambient pressure to stationary phase are composed of primarily saturated, branched fatty acids (67% of total), consistent with lipid content in a wide range of *Bacillus* spp. (50) and consisting of the major fatty acids i15:0, ai15:0, i16:0, n16:0, i17:0, ai17:0 and n18:0, composing 83% of total fatty acids, with a larger percentage of i16:0, n16:0, and n18:0, and a lower percentage of i15:0 compared to its closest relatives (via 16S rRNA identity) (Supplemental Table 5) (51–53).

Stationary phase cultures grown anaerobically under 1 atm and 100 atm of N_2_ or CO_2_ generally have a lower proportion of branched fatty acids (both iso and anteiso forms) relative to aerobically grown cultures, with branched fatty acids composing 29% of total lipids under 100 atm CO_2_, 42% under 100 atm N_2_, 59% under 1 atm CO_2_, and 58% under 1 atm N_2_ (Fig. 1). The branched fatty acid i16:0 varies significantly among the five headspace and pressure combinations with greater abundance under aerobic conditions than all anaerobic conditions (ANOVA p< 0.0003; F-ratio = 15.8, Tukey’s post-hoc test alpha > 0.05) (Supplemental Fig. 4). In contrast, the straight fatty acid, n16:0 is more abundant under all anaerobic conditions, but this difference only met significance between aerobic, ambient pressure and the two high pressure conditions (ANOVA p < 0.02, F-ratio = 5.6, Tukey’s post-hoc test alpha > 0.05).

**Figure 1.**
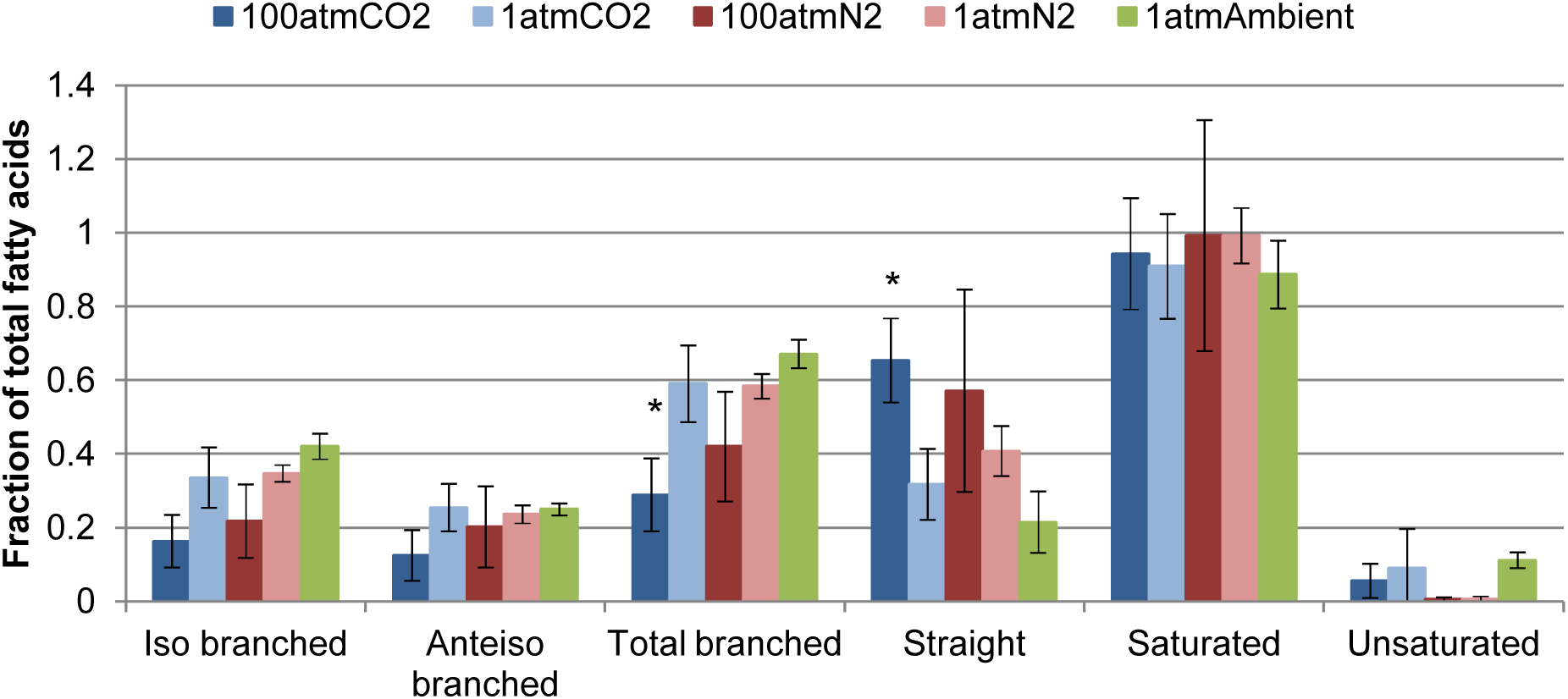
Major lipid classes of MITOT1 sampled in stationary phase under 5 headspace and pressure conditions. Significance described in the text is denoted with (*). Significantly more saturated, normal fatty acids were present in scCO_2_-grown cultures when compared to both 1 atm CO_2_ and 1 atm ambient cultures (t-test p=0.04, p=0.02, respectively). Iso and anteiso branched fatty acids are summed to make up ‘Total branched’ fatty acids. Total branched and normal fatty acids are summed to make up saturated fatty acids. Saturated and unsaturated fatty acids sum to 1 for each sample.

Among the four anaerobic conditions, significant variation was observed for branched fatty acid i15:0 with decreased abundance in 100 atm-incubated cultures relative to those incubated at 1 atm (Two-factor ANOVA p = 0.04; F-ratio=5.6). Significantly more saturated, straight fatty acids were present in scCO_2_ grown biomass when compared to both 1 atm CO_2_ and 1 atm ambient atmosphere grown samples (t-test p=0.04, p=0.02, respectively) (Fig. 1). Fatty acid n16:0 is also significantly elevated under 100 atm CO_2_ relative to 1 atm CO_2_ (t-test p=0.01). However, between all anaerobically grown samples, most individual fatty acids do not show significant differences with respect to headspace and/or pressure by ANOVA or pairwise t-test.

Cultures incubated under 100 atm CO_2_ are associated with the longest average acyl chain lengths (Fig. 2) reaching statistical significance for comparison to 1 atm N_2_ and 1 atm ambient headspace samples (t-test p < 0.04). Several unsaturated fatty acids were only detected in aerobic samples (two unsaturated n16 and two unsaturated n17 fatty acids). However, no significant differences in total unsaturated fatty acids were observed between the different anaerobic headspace and pressure combinations, by either 2-factor ANOVA or pairwise t-tests. We did not find any evidence of hopanoid production (presence of m/z 191 ion with spectral comparison to the NIST database) for strain MITOT1. The same lipid analyses with a different scCO_2_-tolerant *Bacillus* strain (*B. cereus* MIT0214) (3, 40) grown under identical headspace and pressure combinations yielded qualitatively similar results of longer chain lengths and decreased proportion of branching fatty acids under CO_2_ headspaces (Supplemental Fig. 5A-C).

**Figure 2.**
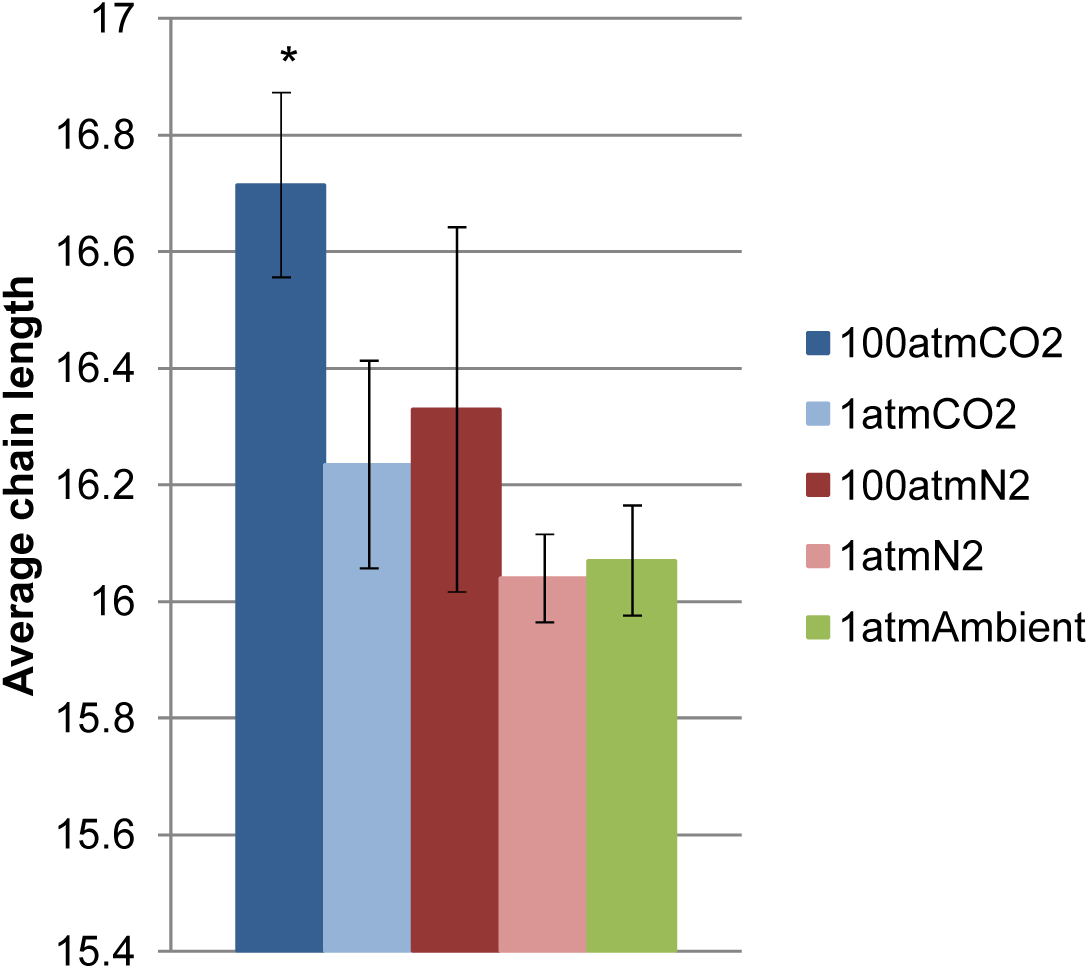
Average fatty acid chain lengths of MITOT1 sampled in stationary phase under 5 headspaces and pressure conditions. Significance described in the text is denoted with (*). 100 atm CO_2_ chain lengths are significantly greater than 1 atm N_2_ and 1 atm ambient samples (t-test p<0.04).

### Proteomes of *B. subterraneus* MITOT1 vary by headspace composition and pressure/incubation conditions

Proteomes from *B. subterraneus* MITOT1 cultures incubated under 1 or 100 atm of CO_2_ or N_2_ resulted in observation of 623 distinct proteins, corresponding to 15% of total proteins predicted by RAST annotation of the genome sequence (40)(Supplemental Fig. 6). These protein counts are similar to those recovered from other *Bacillus* proteomes (within a factor of 2) (54, 55), despite limitations due to low biomass densities obtained from anaerobic cultivation.

To determine whether different headspace and pressure conditions resulted in different proteome profiles, we performed clustering and principal component analysis (PCA) on MITOT1 proteomes. Significant differences were observed between the two batches of proteomes (Supplemental Fig. 6, 7), thus further multivariate analysis was restricted to the 8 proteomes prepared by identical methods (N=2 per condition). These 8 proteomes display high correlation (Spearman R > 0.65), with the highest correlation between duplicates of 1 atm N_2_ and 1 atm CO_2_ (Spearman R > 0.80). The first two principal component axes (Fig. 3) accounted for 79.7% of proteome variation and resolved samples by conditions, with N_2_ and CO_2_ headspace samples separated along PC1 (40.4% variation), and 1 atm and 100 atm incubated cultures separated along PC2 (39.2% variation).

**Figure 3.**
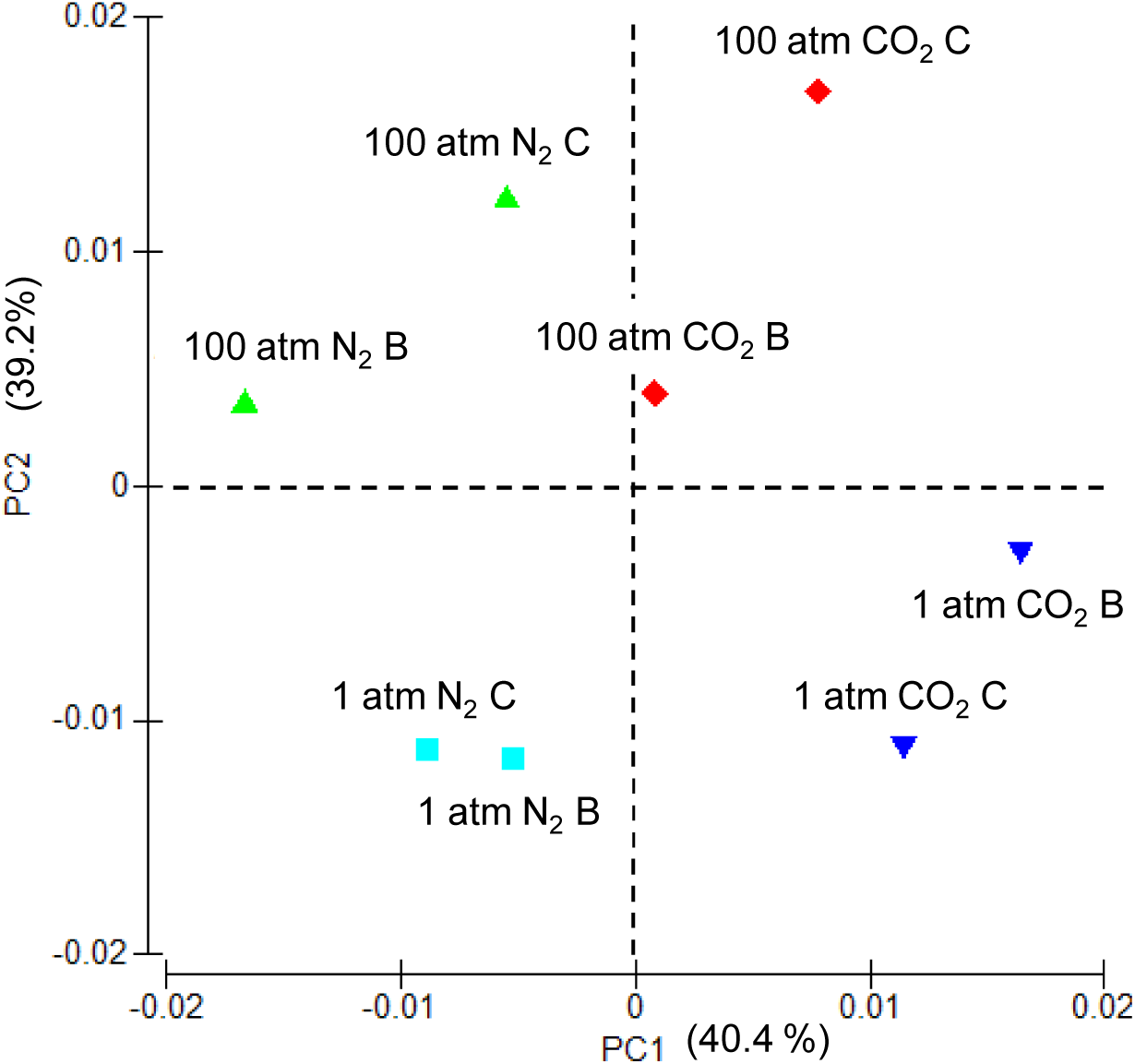
Principal component analysis of duplicate MITOT1 proteomes. Samples separate by headspace along principal component 1, and by pressure along principal component 2, with the first two principal components explaining 79.6% of variability.

### MITOT1 proteomes contain similar profiles of highly represented proteins across conditions

The most abundant proteins in MITOT1 proteomes are consistent with highly expressed proteins present in other *Bacillus* spp. across multiple growth phases (54–56) including predicted functions of translation (e.g. translation elongation factor Tu and G), energy generation (e.g. ATP synthase, and citric acid cycle proteins such as aconitase hydratase and succinyl-CoA ligase), and general stress response (e.g. heat shock protein 60 GroEL and chaperone protein DnaK) (Supplemental Tables 1 A-D).

Among the most highly expressed proteins across conditions, is a hypothetical protein (Fig_2630) predicted to participate in S-layer formation by top BLASTx result (44% identity with 98% coverage to a *B. infantis* NRRL B-14911 S-layer protein) and structural prediction with Phyre2 (48) (42% of the protein predicted) (Supplemental Fig. 8 A). Expression varied from 1.7 – 24.2% of the proteome under CO_2_ and from below detection to 3.6% of the proteome under N_2_ headspaces (Supplemental Fig. 8 B). A second hypothetical protein with structural homology to S-layer proteins (Fig_2635; 27% of the protein predicted) was observed, representing 0.13% - 0.66% of the proteome (Supplemental Table 4). Previous studies document upregulation of S-layer proteins under elevated CO_2_ and low pH conditions (57–59), however, we find predicted S-layer proteins among the most highly represented in the proteome across all conditions and do not vary significantly across anaerobic headspace or pressure (Kruskal-Wallis p > 0.05).

### Proteomes of MITOT1 cultures are consistent with stationary phase growth

Since reduced ribosomal protein expression (5-10 fold) has been associated with stationary phase growth (60) and reduced environmental activity (61, 62), we examined the abundance of total and individual ribosomal proteins in MITOT1 proteomes derived from cultures incubated under different headspace and pressures. Overall, we noted the proportion of ribosomal proteins varied by <2-fold in MITOT1 proteomes (Supplemental Fig 9. A) where high pressure incubated samples were associated with higher ribosomal levels (GSEA p=0.006, FDR<0.25) (Supplemental Fig. 9 B). Removal of ribosomal proteins from the dataset did not significantly change clustering results (Supplemental Fig. 10). Ribosomal proteins L7 and L12, previously shown to be induced during stationary phase growth (63) were observed in all MITOT1 proteomes, and did not vary significantly by headspace or pressure (Kruskal-Wallis p=0.78).

The abundance of several additional predicted proteins suggest proteomes were collected within the targeted stationary phase (Supplemental Fig. 11). Observation of proteins predicted to participate in acetoin metabolism (Fig_989, Fig_1512 and Fig_3152) are consistent with their previous observation in stationary-phase (but not exponential-phase) proteomes of *B. thuringiensis* (55) and may reflect the role of acetoin as a secondary carbon source during stationary phase growth of multiple bacterial types (64). In addition, the MITOT1 proteomes revealed a carbon starvation protein (Fig_2079), the cell division protein, FtsH (Fig_3548) and two cold shock proteins (Fig_446 and Fig_3723) which have also previously been associated with proteomes of stationary phase cultures (65, 66). Although many *Bacillus* species undergo sporulation during stationary phase, relatively few predicted sporulation proteins were detected, under all anaerobic conditions, consistent with microscopic observation of cells with primarily vegetative rather than spore morphology during direct counts (Supplemental Fig. 12) and also noted elsewhere (Peet et al, (3)).

### Glycine cleavage system is enriched in CO_2_ headspaces

A total of 476 proteins (of 623 total) were annotated with KEGG and corresponded to 43 pathways which were analyzed by GSEA. Six of 43 pathways were significantly enriched with respect to either headspace or pressure (p<0.05). We focused on differences between gas headspace to control for the potential effects of variability in culture age within stationary phase, (i.e. pooling 1 atm and 100 atm samples from each headspace). The pathway for glycine, serine, and threonine metabolism was enriched under CO_2_ headspace, while no pathways were significantly enriched under N_2_ (Supplemental Table 2). Five of the proteins in the pathway for glycine, serine, and threonine metabolism (Figure 4) comprise the glycine cleavage system, which is involved in glycine and serine catabolism or synthesis. The protein most highly correlated with CO_2_ samples through PLS-DA is glycine dehydrogenase (decarboxylating) enzyme, which is noteworthy as it can either produce or consume CO_2_ depending on which direction the reaction proceeds, and has been previously shown to be upregulated in acid stressed *E. coli* (67). All 5 proteins involved in the glycine cleavage system were positively correlated with CO_2_ headspaces.

**Figure 4.**
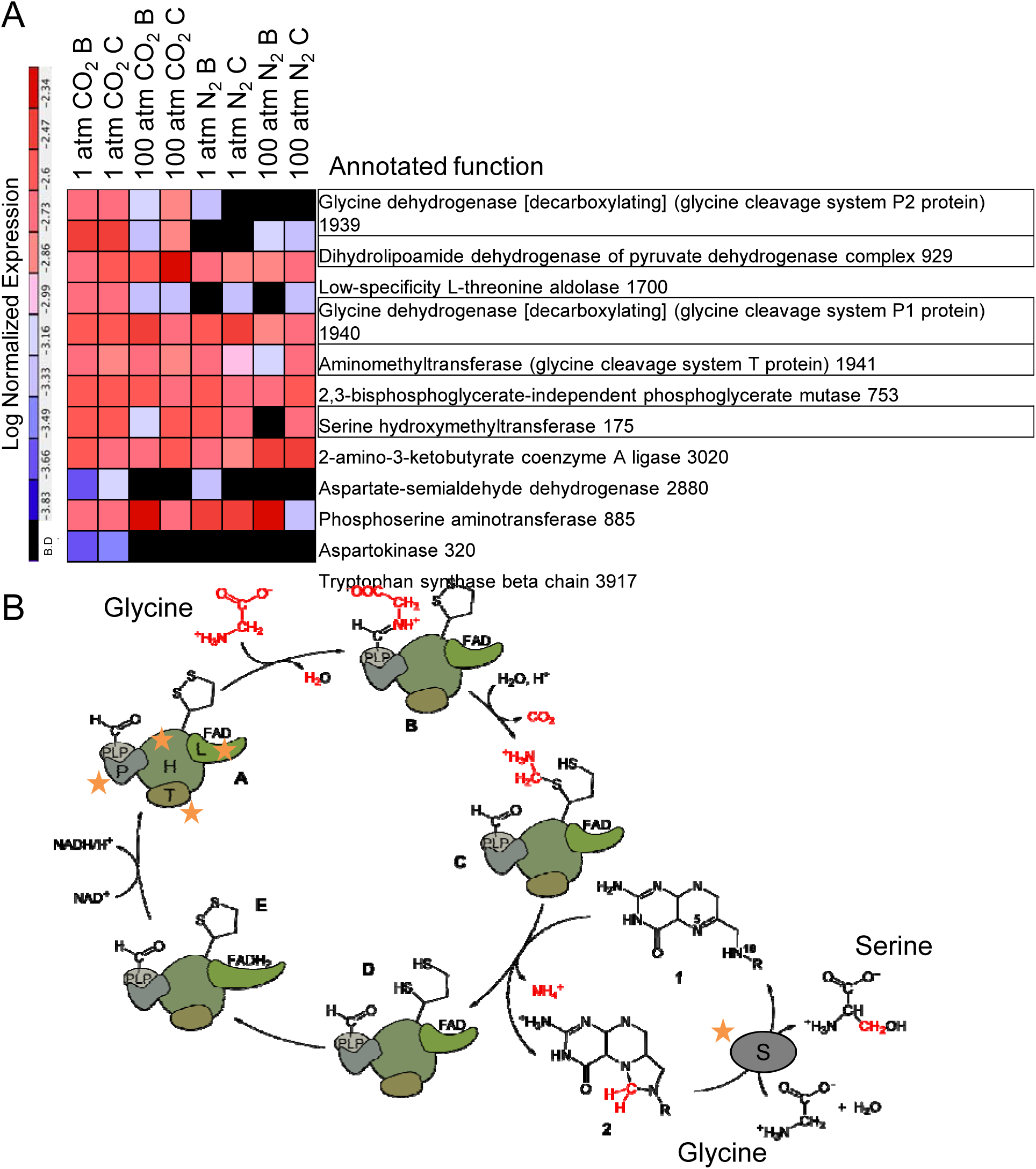
Enrichment of glycine, serine and threonine metabolism under CO_2_. (A) Heatmap of spectrum counts for proteins in the KEGG gene set for glycine, serine and threonine metabolism. Enrichment under CO_2_ headspace samples indicated via GSEA (nominal p = 0.018). Proteins from the Glycine cleavage system are outlined and are all positively correlated with a CO_2_ headspace, (PLS-DA p(corr) of 0.98 for Glycine dehydrogenase P2 protein [H], 0.90 for Dihydrolipoamide dehydrogenase [L], 0.85 for Glycine dehydrogenase P1 protein [P], 0.42 for Aminomethyltransferase [T], and 0.40 for Serine hydroxymethyltransferase [S]). The final number on each protein annotation is the RAST annotation fig number. (B) The Glycine cleavage system for glycine metabolism can either produce or consume CO_2_. Letter codes on diagram correspond to names followed by code in square bracket given in part A. Modified with permission: (90) (https://en.wikipedia.org/wiki/Glycine_cleavage_system#/media/File:Glycine_decarboxylase_complex.svg).

PLS-DA revealed that additional proteins involved in amino acid metabolism, including deblocking aminopeptidase and arginase, were highly correlated with CO_2_ grown cultures (Supplemental Table 3 A). Deblocking aminopeptidase is potentially significant as it is involved in amino acid metabolism and it is upregulated during heat and oxidative stress (68). Arginase is notable as it is involved in the *H. pylori* acid stress response by producing urea from arginine (32). Additional predicted proteins with expression that is highly correlated with CO_2_ from PLS-DA include stress response and respiratory proteins (Supplemental Table 3 A).

Predicted proteins with differential expression between N_2_ and CO_2_ (Kruskal-Wallis test, p<0.05) include several that are involved in energy generation pathways (Supplemental Fig. 13 E-G). Aconitase hydratase is a TCA cycle protein that is abundant in all conditions, although it is significantly higher under CO_2_ headspaces. However, other TCA cycle enzymes do not show significantly different expression under CO_2_ or N_2_. An electron transfer flavoprotein, beta subunit is significantly reduced under CO_2_, with another electron transfer flavoprotein showing similar patterns of expression, but not significantly so. Similarly, a tungsten-containing aldehyde:ferredoxin oxidoreductase implicated in carbon utilization (69) showed significantly lower expression under CO_2_.

Additional predicted proteins with differential expression between N_2_ and CO_2_ conditions include proteins involved in stress response and/or protein folding (i.e. Chaperone Dnak, ATP-dependent Clp protease ATP-binding subunit ClpX, ATP-dependent hsl protease ATP-binding subunit, and a Universal stress protein). These all show significantly decreased expression under CO_2_ (Kruskal-Wallis test, p<0.05) (Supplemental Fig. 13 A-D).

### Discussion

Much of the current literature on microbial responses to supercritical CO_2_ involves shocking vegetative cells with rapid increases in headspace CO_2_ content, leading to increased pressure, and acidity and a loss in cell viability (4, 5, 8, 9, 11, 70). In contrast, the model system for scCO_2_ exposure employed in this study allows examination of living cells where cultures are inoculation with spores, germination occurs after scCO_2_ addition, and the resulting protein and lipid profiles reflect acclimated growth. In this study we have examined, for the first time, how proteins and lipids vary in different headspace and pressure/incubation conditions to gain insight into how cells acclimate to growth under scCO_2_. To target similar growth phase in cultures incubated across conditions with variable pressure and headspace we restricted this study to cultures predicted to be in stationary phase based on growth curves conducted at 1 atm or the growth-frequency based logistic regression analysis described in Peet et al (3), for cultures grown at 100 atm. Indeed, observed cell density and protein expression across headspace and pressure/incubation conditions are consistent with stationary phase growth. Congruence of highly expressed proteins in scCO_2_-exposed proteomes with other *Bacillus* proteomes from stationary phase cultures (55, 56, 71) is notable as it suggests that *B. subterraneus* MITOT1 can acclimate and maintain housekeeping metabolic processes under a scCO_2_ headspace.

Analysis of acclimation of *B. subterraneus* MITOT1 to scCO_2_ through lipid and proteome profiling supports the hypothesis that resistance to CO_2_ stress is similar to an acid stress response, as the high concentration of CO_2_ (associated with a supercritical headspace) results in a significant pH reduction in the growth media (3). Analysis of lipids suggests a reduced proportion of branched fatty acyl lipids and increased average acyl chain lengths under scCO_2_ in strain MITOT1 that may increase membrane rigidity. These observations are supported by qualitatively similar trends observed in *B. cereus* MIT0214, notably longest average lipid chain lengths under 100 atm CO_2_ (Supplemental Fig. 5) (72). These lipid changes observed in cells grown under scCO_2_ suggests that modulation of membranes to be less fluid may be an important acclimation mechanism in response to the membrane permeabilizing effects of scCO_2_ and is similar to changes observed in acid stressed (22) and solvent stressed bacteria (21).

While our data do not reveal significant differences in the abundance of predicted S-layer proteins between N_2_ and CO_2_, existing literature from *Bacillus* strains indicate that S-layers may facilitate acclimation to acid and CO_2_ stress (57, 58, 73, 74). S-layers also play a role in cell adhesion, virulence, and membrane stability as they form a protein lattice on the cell surface (59, 75). Given the stresses that scCO_2_ imposes (e.g. cytoplasm acidification and membrane permeabilization), and the changes observed in membrane lipids, we hypothesize that universally high S-layer production may predispose MITOT1 to survive and grow under diverse environmental stresses, including those associated with scCO_2_.

Principal component and clustering analyses indicated that both headspace and pressure help explain variability in the MITOT1 proteomes, with samples separating by headspace along the first principal component and by pressure along the second principal component (Figure 3). Interestingly, the protein profiles of cells grown under scCO_2_ appears to be intermediate between low pressure CO_2_ and high pressure N_2_, consistent with our hypothesis that acidity and pressure may have some opposing effects.

The protein most highly correlated with the CO_2_ headspace condition was glycine dehydrogenase, a component of the glycine cleavage system, which has been demonstrated to be upregulated in acid stressed *E. coli* (67). Enrichment of amino acid metabolism in CO_2_ conditions, in particular five proteins mediating the glycine cleavage system, supports the hypothesis that this system may play a role in CO_2_ acclimation. Two other proteins enriched under CO_2_ conditions, arginase and deblocking aminopeptidase, have been previously shown to be pH or stress-responsive. Arginase is associated with pH neutralization (32) and deblocking aminopeptidase is upregulated during heat and oxidative stress (68). These amino acid metabolic proteins represent future targets for understanding how MITOT1 responds to elevated CO_2_ stress, as amino acid metabolizing pathways are involved in acid neutralization through production of neutralizing compounds (27, 76) and consumption of protons (28), while various amino acids have been documented as compatible solutes for osmotic regulation (77).

The stationary phase cultures examined in this study reflect cells that are no longer growing, and these share some similarities with natural settings where a large portion of cells are static i.e. in dormant forms such as spores or in very slow growing states (78, 79). Bacteria grown under continuously-diluted culture conditions mimicking static conditions reveal lower expression of DNA and protein repair than stationary phase cultures where toxic end products may accumulate and damage cells (80). Both non-static and stationary-phase populations must downregulate ribosomal proteins and energy generation pathways as growth rate slows, suggesting that stationary phase cultures can approximate some aspects of static populations (78–80).

While doubling times for static subsurface microbial populations are estimated to exceed hundreds of years (81), these populations can influence the biogeochemistry of the subsurface and may respond to perturbations in their environment. Stimulated growth of static microbial populations, such as spores, may be relevant in the context of various subsurface industrial activities including enhanced oil recovery, hydraulic fracturing, and geologic carbon sequestration. For example, active subsurface communities can affect variables such as porosity and permeability through mineral dissolution and nucleation (2, 82, 83). Additionally, cells in static or stationary phases may be more resistant to some stresses (8, 84, 85) and thus more likely to survive and grow after a perturbation in the subsurface (such as scCO_2_ exposure during geologic carbon sequestration (GCS)). Indeed, it has been inferred that microbial communities in the deep subsurface may be acclimating to influxes of scCO_2_ during GCS through changes in community composition (86, 87).

The demonstration of microbial growth under a scCO_2_ headspace calls into question the use of scCO_2_ as a food and medical sterilizing agent where spores may be present, but it is encouraging for the development of applications involving scCO_2_ including bioengineering in GCS and biocatalysis under scCO_2_. Biocatalysis under scCO_2_ is currently conducted with enzymes or inactivated cells (88). Biphasic reactors containing scCO_2_ and an aqueous (or other solvent) phase have been explored for extraction of various biologically produced compounds that partition into the scCO_2_ phase (1, 89). This study provides insights into how live cells maintain biocompatibility with scCO_2_ through shifts in lipid composition and protein expression, and provides candidate targets to improve the growth of non-biocompatible strains. Membrane lipid modifications to create a more rigid membrane and activity of the glycine cleavage system may help cells acclimate to scCO_2_. In addition, we hypothesize that introduction of cells as spores to scCO_2_ systems, followed by germination and growth of acclimated cells may help *Bacillus* spp. tolerate the complex stresses associated with scCO_2_. New opportunities for biotechnology development in biofuels, pharmaceuticals, and microbially-enhanced GCS will be possible with microbes that are able to grow in an aqueous phase in contact with scCO_2_. For any of these applications to be realized, the continued investigation and development of supercritical CO_2_ tolerant microorganisms like *B. subterraneus* MITOT1 is crucial.

## Acknowledgements

We would like to thank Kelden Pehr and Florence Schubotz for assistance with lipid analysis. Funding was provided to JRT by the Department of Energy under awards DE-FE0002128, DE-SC0012555 and by the MIT Energy Initiative. Lipid analyses were supported by a grant (NNA13AA90A) from the NASA Astrobiology Institute to RES. The proteomics experiments and bioinformatics were carried out in the MIT Center for Environmental Health Sciences Bioanalytical Core Facility, supported by NIEHS Grant #P30-ES002109.Disclaimer: “This publication was prepared as an account of work sponsored by an agency of the United States Government. Neither the United States Government nor any agency thereof, nor any of their employees, makes any warranty, express or implied, or assumes any legal liability or responsibility for the accuracy, completeness, or usefulness of any information, apparatus, product, or process disclosed, or represents that its use would not infringe privately owned rights. Reference herein to any specific commercial product, process, or service by trade name, trademark, manufacturer, or otherwise does not necessarily constitute or imply its endorsement, recommendation, or favoring by the United States Government or any agency thereof. The views and opinions of authors expressed herein do not necessarily state or reflect those of the United States Government or any agency thereof.”

The authors declare no conflict of interest with this work.

